# Potential mechanisms and effects of melatonin-regulated Nrf2/HO-1 pathway on acute lung injury due to formaldehyde exposure

**DOI:** 10.1101/2024.12.17.629008

**Authors:** Bihong Wang, Jianguo Lv, Miao Xu, Dewei Chang, Zhe Wu, Yanling Sun

## Abstract

Acute lung injury is a topic of great interest in critical care medicine due to its high mortality rates. The lungs are the immediate target organ for formaldehyde inhalation damage. Lung damage and fibrosis are the most important outcomes of severe and acute lung disease and pose a serious threat to human health. Melatonin (MT), a natural bioactive compound with anti-inflammatory and antioxidant properties, However, it is not clear whether MT can prevent FA-induced acute lung injury (ALI). Therefore, in this study, we aimed to evaluate the protective effects of MT and the potential mechanisms against FA-induced ALI. An environmental exposure bin was used to inhale 3 mg·m3 FA-induced ALI, which was given intraperitoneally with different doses of MT (5/10/20 mg/kg) after successful modeling. In addition, rats were treated with Nrf2 inhibitor (ML385) to validate the signaling pathway. Lung function was measured, histopathological/morphological changes in lung tissue were assessed, and inflammatory expression and oxidation levels in lung tissue were detected. We observed that MT greatly alleviated the lung dysfunction, pathological lung injury, pulmonary edema and inflammatory response after successful modeling of FA. In additional, MT played a role in modulating the Nrf2/HO-1 signaling pathway, which effectively inhibit oxidative stress caused by FA-induced lung tissue injure. Moreover, we found that activation of the NF-κB pathway is associated with inflammation caused by this injury. Overall, our data suggest that MT inhibits the expression of oxidative stress and inflammation in lung tissue through the institutional or Nrf2/HO-1 pathway, alleviating FA-induced ALI.

## 1. Introduction

Acute lung injury (ALI) is an early lesion of ARDS, ALI and more severe ARDS as common mortality and life-threatening lung diseases. Although some progress has been made in the diagnosis and treatment of ALI/ARDS, the pathogenesis is very complex due to its many causative factors.^1^ There are still no effective therapeutic measures, which allows the mortality rate of the disease to remain as high as 40% and seriously affects the prognosis of critically ill patients.^2^ The protein-rich edematous fluid in ALI/ARDS is associated with large numbers of neutrophils, pro-inflammatory cytokines and cytokines, proteases and oxidants.^3^

Formaldehyde (FA) is widely used in modern industry and is a widespread environmental and occupational pollutant, and among all known health effects of FA, lung injury is one of the most serious risks. Millions of people worldwide are exposed to FA every day.^4^ Studies have shown that FA leads to ALI through reduced transalveolar Na^+^ transport, reduced human epithelial sodium channel activity and enhanced membrane depolarization, and increased ROS production.^5–6^ ROS upregulate inflammatory cytokines and perpetuate malignancy by recruiting more inflammatory cells to perpetuate the vicious cycle, ultimately leading to severe tissue damage.^7^ Addressing inflammation and oxidative stress, which can lead to lung injury, is a desirable goal in the treatment of FA-induced ALI. Currently, there are no effective therapeutic agents and preventive strategies in clinical practice. And multiple drug candidates with novel and unique mechanisms of action are needed.

Melatonin (MT) has been reported to play a key role in various physiological activities, including the regulation of circadian rhythms, immune responses, oxidative processes, apoptosis or mitochondrial homeostasis,^8^ and its most prominent pharmacological effects are the scavenging of free radicals and the inhibition of inflammatory responses.^9^ Recent studies have shown that MT is an important antioxidant and anti-inflammatory carrier that plays a crucial role in alleviating oxidative stress and overproduction of pro-inflammatory cytokines and chemokines in lung tissues,^10^ while the beneficial effect of MT is associated with nuclear factor erythroid 2-related factor 2 (Nrf2) activation.^11^

Nuclear factor E2-related factor 2 (Nrf2) is a major transcriptional regulator that ensures the protection of a large number of tissues and cells from ROS-mediated induction due to its various antioxidants and phase II detoxification enzymes,^12^ ^12^activates the transcription of antioxidant genes and is also involved in the regulation of cell proliferation and inflammatory gene expression.^13^ Large amounts of ROS activate tyrosine kinases to dissociate the Nrf2: Keap1 complex, nuclear import of Nrf2 and coordinated activation of cytoprotective gene expression.^14^ At the same time, oxidative stress activates cellular NF-κB inflammatory signaling and leads to chronic inflammation.^15^ Nrf2 promotes anti-inflammatory processes through cross-talk with the NF-κB pathway.^16^

Considering the ubiquity of FA in urban areas due to environmental pollution, it is important to explore effective strategies to stop the health hazards associated with FA. Based on this, we hypothesized that MT could exert a protective effect on FA-induced ALI through activation of Nrf2. Therefore, our study aimed to investigate the protective role of MT in FA-induced ALI and to explore the underlying molecular mechanisms.

## 2. Materials and Methods

### 2.1 Animals and treatment

A total of 60 famale Wistar rat weighing 130-150g (5-6 weeks old) were purchased from Hubei Province Experimental Animal Center (Wuhan, China). All animals were housed in a 12 h light/dark circumstance with food and water ad libitum. All experimental procedures were performed according to the local and international guidelines on the ethical use of animals, and all efforts were made to minimize the number of animals used and their sufferings. Ethics approval was obtained from the Laboratory Animal Ethics Committee of Hubei University of Science and Technology (2019-03-021). After a week of acclimatization feeding, we randomly divided 60 female Wistar rats into six groups: Control, FA, FA+MT5mg/kg, FA+MT10mg/kg, FA+MT20mg/kg, FA+MT10mg/kg+ML385 (APEXBIO; B8300; America), with 10 rats in each group. All groups, except for the Control group, were exposed to FA (aladdin; F11941; China) 3mg/m3^17–18^ through intranasal inhalation for 4 hours per day for 21 days (IES-NI; China). Following this, they were treated with intraperitoneal injections of MT (aladdin; M18674; China) at doses of 5/10/20mg/kg ^19–20^ for 14 days, while continuing to be exposed to FA. The Control group, on the other hand, was injected intraperitoneally with an equal volume of 0.9% sodium chloride solution. Refer to (Fig.1) for further detail.

**Fig. 1.**
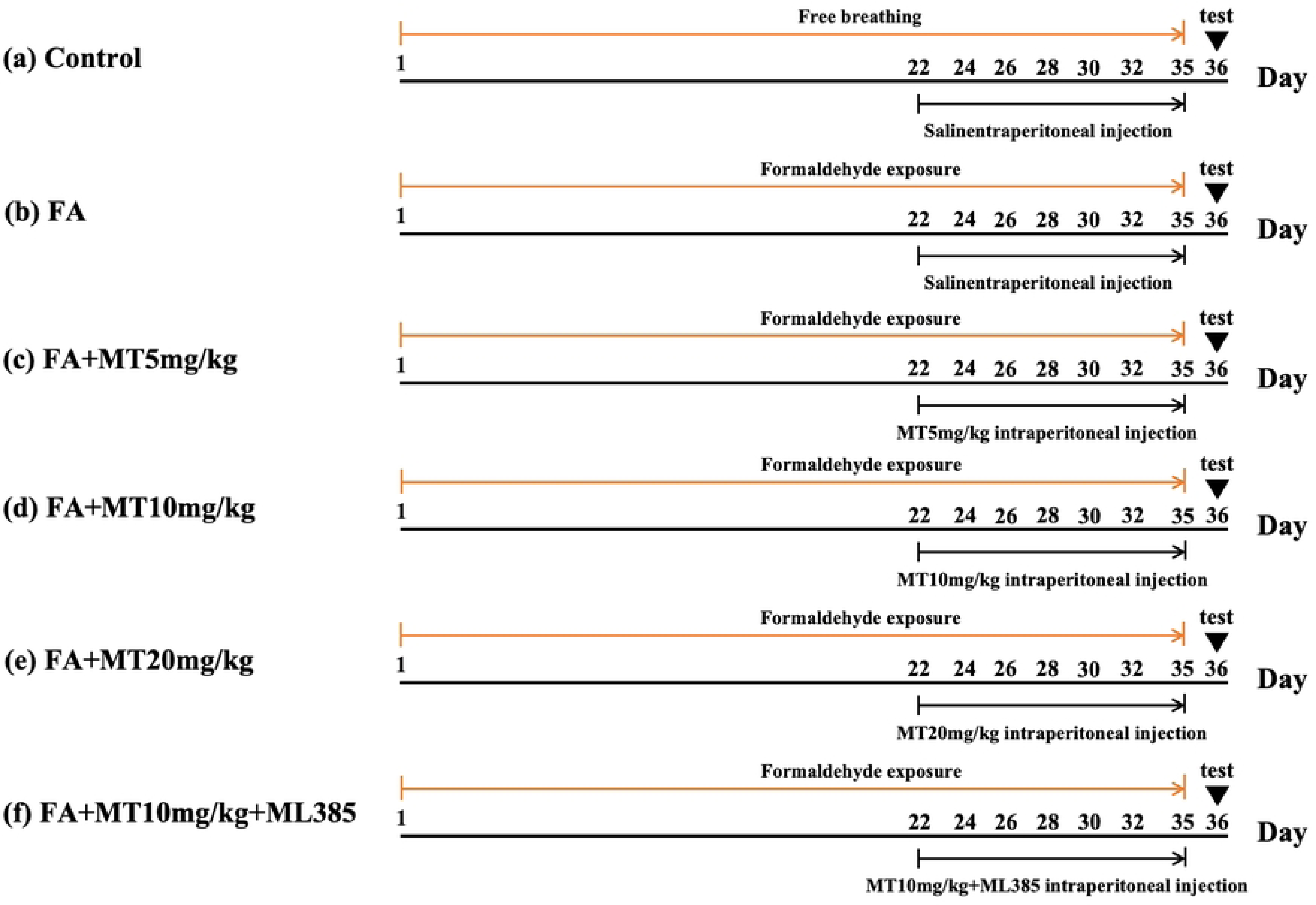
Schematic diagram of the experimental procedures

### 2.2 Measurement of Airway hyperresponsiveness (AHR)

According to the manufacturer’s instructions of the AniRes2005 lung function system (Bestlab, version 2.0, China), Rats were anesthetized by intraperitoneal injection of 1% pentobarbital sodium (Urchem, China). The respiratory rate was pre-set at 90/min, and the time ratio of expiration/inspiration was 20: 10. AHR was assessed by the indexes of Re, Ri, and the minimum value of Cldyn. Ri and Re R-areas, the graph area between the peak value and baseline, and the valley of Cldyn were recorded for further analysis.

### 2.3 Lung Histological Assay

After ventilator testing, lung tissues were removed and fixed in 4% paraformaldehyde (PFA, 0.1 M phosphate buffer, pH 7.4) at 4°C for 12 h. The tissues were then embedded in paraffin wax and cut into 4-μm sections with a microtome (RM 2165; Leica Microsystems GmbH). Sections were stained with haematoxylin and hemoglobin (H&E), periodic acid-Schiff (PAS), and Masson’s trichrome to assess the level of inflammation or fibers in the lungs (Solarbio; G1120; G1285; G1346; China). Briefly, the sections were deparaffinized with xylene, 100% ethanol, 90% ethanol, and 70% ethanol, and then treated with staining solutions, stained, and sealed with neutral resin, and then visualized with a fluorescence microscope (Olympus IX73; Olympus). The degree of alveolar edema, intra-alveolar congestion, interstitial edema, and intra-alveolar congestion were assessed and scored separately in this study. Score of 0 indicates no change or very slight change, 1 indicates slight change, 2 indicates moderate change, 3 indicates severe change, and 4 indicates very severe change. The scores of these four items were averaged to obtain the H&E staining lung injury score.

### 2.4 Lung wet/dry (W/D) weight measurement

The body mass was first weighed, and then the anterior lobe of the right lung was extracted to calculate the lung coefficient, which is the ratio of lung weight to body mass. To determine the surface water content of the lung tissue, filter paper was used to absorb the wet weight, which was then recorded. The tissue was then dried in a constant temperature oven at 70℃ for 48 hours and weighed again to obtain the dry weight, which was used to calculate the W/D ratio (wet weight divided by dry weight). The lung water content was calculated to reflect the degree of pulmonary edema.

### 2.5 Molecular docking

The X-ray crystal structure of Nrf2 was obtained from the Protein Data Bank (PDB ID: 1X2R https://www.rcsb.org/). The structure of MT was downloaded from the PubChem database (https://www.pubchem.ncbi.nlm.nih.gov/compound) and optimized using ChemBio3D Ultra 14.0 software (PerkinElmer Informatics). Auto Dock Vina 1.1.2 software (Center for Computational Structural Biology) was used to dock conformation between Nrf2 and MT. PyMOL 2.2.3 was used to visualize the conformation.

### 2.6 Immunohistochemistry (IHC)

lung tissue sections were dewaxed, conducted to antigen retrieval (Beyotime Biotech; P0083; China), treated with 3% hydrogen peroxide for 10 min, closed with 10% goat serum closure solution (Concentrated SABC-POD Rabbit IgG Kit; Boster BiolTech; SA2002; China) for 1 h, and then incubated with primary antibody overnight at 4°C, After the primary antibody was applied, the sample was incubated with secondary antibodies at room temperature for 1h. The peroxidase present in the secondary antibodies was utilized to oxidize the DAB, resulting in the formation of a brownish-yellow precipitate with the DAB chromogenic solution. Following this, the nuclei were stained blue with hematoxylin and observed under a fluorescence microscope (Olympus IX73; Olympus Corporation). The fluorescence intensities were analyzed using ImageJ 1.51j8 (National Institutes of Health). The following primary antibodies were used: anti-Nrf2 (1: 100; proteintech; 16396-1-AP; America), anti-HO-1 (1: 100; proteintech; 10701-1-AP; America) and anti-NF-κB (1: 100; proteintech; 10745-1-AP; America).

### 2.7 Western blotting

The study utilized the right posterior lobe of the animal’s lung, homogenized in RIPA lysis buffer containing protease inhibitors (SEVEN Biotech; SW105; SW107-02; China), centrifugated at 12,000 g, 4°C for 20 min. Then the supernatant was collected, separated on SDS-PAGE, and transferred to 0.22 μm PVDF membranes. Protein concentration was quantified using a BCA analysis kit (Abbkine; KTD3001; America). Then the membranes were blocked with QuickBlockTM Blocking Buffer for Western Blot (Biosharp Life Sciences; BL502A, China), incubated with the appropriate primary antibodies overnight at 4°C. And HRP-conjugated secondary antibodies in TBST (1: 5,000) at room temperature for 1 h. Protein bands were visualized using ECL detection reagent (Abbkine; K22030; America) and detected with an iBright 1500 instrument (Invitrogen; Thermo Fisher Scientific, Inc). The grey values of bands were analyzed using ImageJ 1.51j8 software (National Institutes of Health). β-actin was used as a loading control. The following primary antibodies were used: anti-HO-1 (1: 1000; proteintech; 10701-1-AP; America), anti-Nrf2 (1: 1000; Abbkine; ABP0106; America), anti-NF-κB (1: 1000; proteintech; 10745-1-AP; America) and anti-p-NF-κB (1: 1000; Abbkine; ABP0043; America).

### 2.8 Fluorescence quantitative PCR

The study utilized the right posterior lobe of rat lung, from which RNA was extracted and purified using the Tissue Extraction RNA Kit (Dakewe Biotech; 8034111; China). Reverse transcription was performed using the All-in-one First Strand cDNA Synthesis Kit (SEVEN Biotech; SM31-02; China) to obtain cDNA. The cDNA obtained from the reverse transcription was used as a template and detected using the perfectStartTM Green qpCR SuperMix kit (TransGen Biotech; AQ601-02; China). Table 1 displays all primers used in the study. The mRNA levels were calculated with the 2-△△Ct method and normalized to β-actin. The primer sequences were as follows: Nrf2 5’-TTCAAGCCGATTAGAGG-3’, reverse 5’-TTGCTCCTTGGACATCA-3’; HO-1: forward 5’-GGTCCTGAAGAAGATTGCG-3’, reverse 5’-GATGCTCGGGAAGGTGAA-3’; Keap1: forward 5’-CGCCCTGTGCCTCTATG-3’, reverse 5’-AGGTGCCACTCGTCTCG-3’; and β-actin: forward 5’-CGTTGACATCCGTAAAGACC-3’, reverse 5’-GGAGCCAGGGCAGTAATCT-3’. Enzyme-linked immunosorbent assay (ELISA)

After ventilator assay, the upper end of the tracheal cannula was then inserted using a syringe, and 8 ml of saline was injected into the left lung in three separate doses. The lavage fluid was flushed and recovered three times, and the supernatant was centrifuged for 10 minutes at 12000 r-min-1 under 4 ℃. Finally, the levels of TNF-α, IL-6, and IL-1β were measured using the ELISA kit (Abbkine; America) instructions.

### 2.9 superoxide dismutase (SOD), glutathione (GSH), and 8-Hydroxydeoxyguanosine (8-OHdG) analyses in the lung tissues

To detect oxidative stress indicators, some biomarkers of lipid peroxidation including SOD (Shanghai Biyuntian Biotechnology Institute; S0101S; China), GSH and 8-OHdG (Nanjing Jiancheng Bioengineering Institute; A061-1/H165-1-1; China) were detected using commercial assay kits according to the manufacturer’s instructions.

### 2.10 Statistical analysis

Data were expressed as mean ± standard deviation (SD) and analyzed using Graphpad Prism 9.0. Normal distribution was assessed with the Shapiro-Wilk test, and multiple comparisons were made using one-way ANOVA, followed by Bonferroni test to compare data across multiple groups. Finally, we used a two-way ANOVA with multiple comparison test to analyze the AHR results.

## 3. Results

### 3.1 MT treatment alleviates FA-induced lung function abnormalities

To assess the changes in AHR, we compared airway responses to MeCh in different groups (Fig.2). In all experimental groups, both the expiratory and inspiratory resistance increased with an increase in the MeCh dose, while the trough value of Cldyn decreased. At each point, FA exposure had significant effects on Ri, Re, and Cldyn (p < 0.05 or p < 0.01) in each treatment group. Compared with the FA group, MT could significantly reduce the changes in lung function (p < 0.05 or p < 0.01), mainly including reduced Ri and Re and increased dynamic lung compliance. This indicated that the MT could effectively reduce the changes in lung function induced by FA.

**Fig. 2.**
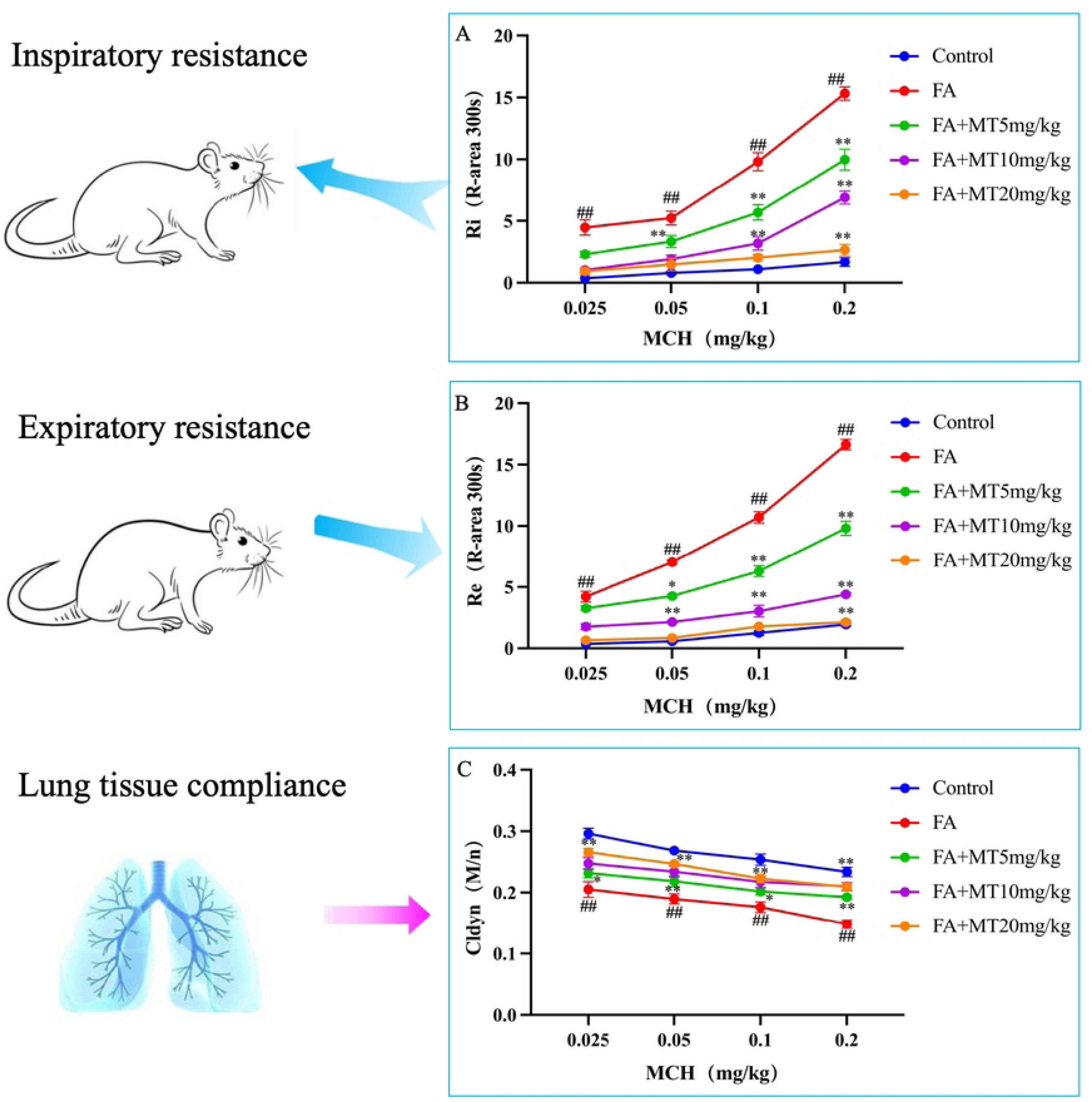
MT treatment alleviates FA-induced lung function abnormalities (A) R-area of Ri, (B) R-area of Re, and (C) peak value of Cldyn at different doses of MeCh Animal groups (in all panels): n = 3 rat per group. (*: p < 0.05, **: p < 0.01, compared with the FA group; ##: p < 0.01, compared with the control group).

### 3.2 MT treatment attenuates FA-induced ALI

In H&E staining lung tissue sections, we observed thickening of alveolar walls, collapse of alveoli, and massive infiltration of inflammatory cells in the FA group (Fig.3A). The W/D ratio and lung factor of lung tissue in FA group were increased, indicating the occurrence of pulmonary edema. MT treatment reduced lung injury scores and greatly attenuated the development of pulmonary edema (Fig.3B-D).

**Fig. 3.**
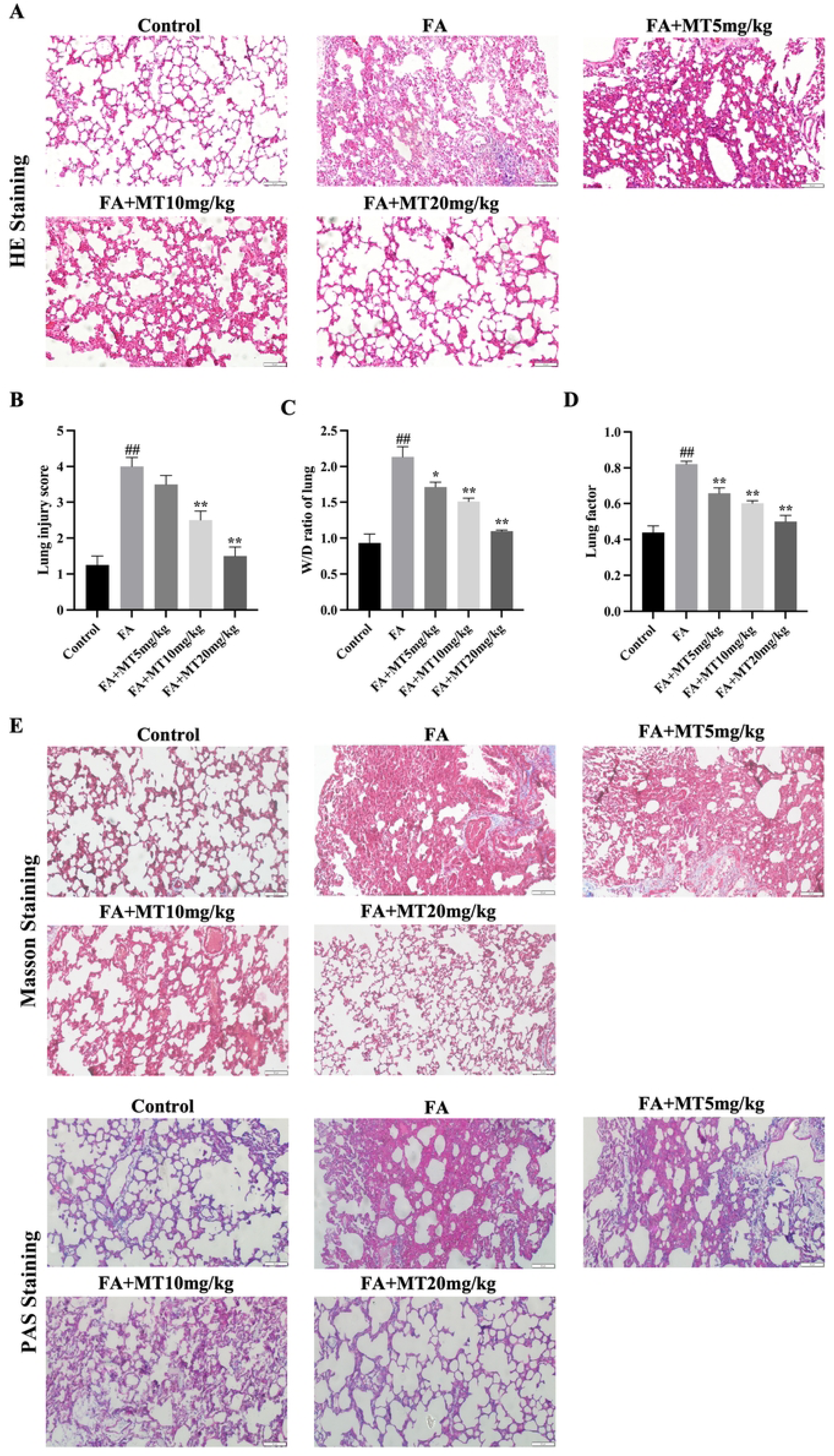
MT treatment attenuates FA-induced ALI (A) Representative H&E staining images of each group of lung tissue sections. Scale bar = 50μm. (B) Quantitative analysis of inflammation score for the H&E staining in each group. (C) Lung wet/dry (W/D) weight ratio. (D) Lung coefficient measurement. (A) (E) Representative Masoon/PAS-stained images of lung tissue sections from various groups. Scale bar = 50μm. Animal groups (in all panels): n = 3 rat per group. (*: p < 0.05, **: p < 0.01, compared with the FA group; #: p < 0.05, ##: p < 0.01, compared with the control group).

Masson staining demonstrated collagen fiber deposition and structural damage, and PAS staining showed significant glycans, which indicated pathological changes such as chronic inflammation in the lung tissue (Fig. 3E). However, treatment with MT reduced these tissue structural abnormalities and inflammatory response induced by FA.

### 3.3 MT antagonizes FA-induced ALI through the Nrf2/HO-1 pathway

A molecular docking assay was performed on the X-ray crystal structures of Nrf2 and the ligand MT (Fig.4A). Auto Dock data showed that MT formed two electrovalent bonds with Nrf2 at residues CLY-367 and ARG-415.The electrovalent bond distances were measured to be 2.2 ng strom between Nrf2 CLY-367 and MT, and 2.3 ng strom between Nrf2 ARG-415 and MT. And the binding affinity was -7.6 kcal/mol. It indicates that Nrf2 has a higher affinity for MT. To investigate the protective mechanism of MT against FA-induced ALI, the level and localization of Nrf2/HO-1 were analyzed by immunohistochemical analysis. The expression level of Nrf2/HO-1 in the lung tissue of rats in FA group was lower than that in the control group, but MT could promote the entry of Nrf2 into the nucleus and up-regulate the expression of Nrf2, suggesting that the protective effect of MT on FA-induced ALI might be related to the up-regulation of Nrf2 (Fig.4B-C). In addition, we demonstrated the protective effect of Nrf2 against FA-induced ALI with Nrf2 inhibitor ML385, which significantly reversed FA-induced oxidative stress in lung tissue. These findings suggest that Nrf2 plays an important role in the development of FA-induced ALI, and that MT may ameliorate FA-induced ALI by activating Nrf2, thereby reducing oxidative stress. The changes of Nrf2/HO-1 expression levels in the results of Western blot and qPCR were consistent with the results of immunohistochemistry (Figure 4D-H), and the mRNA expression level of keap1, the downstream molecule of Nrf2/HO-1, also showed significant changes. (Fig.4I).

**Fig. 4.**
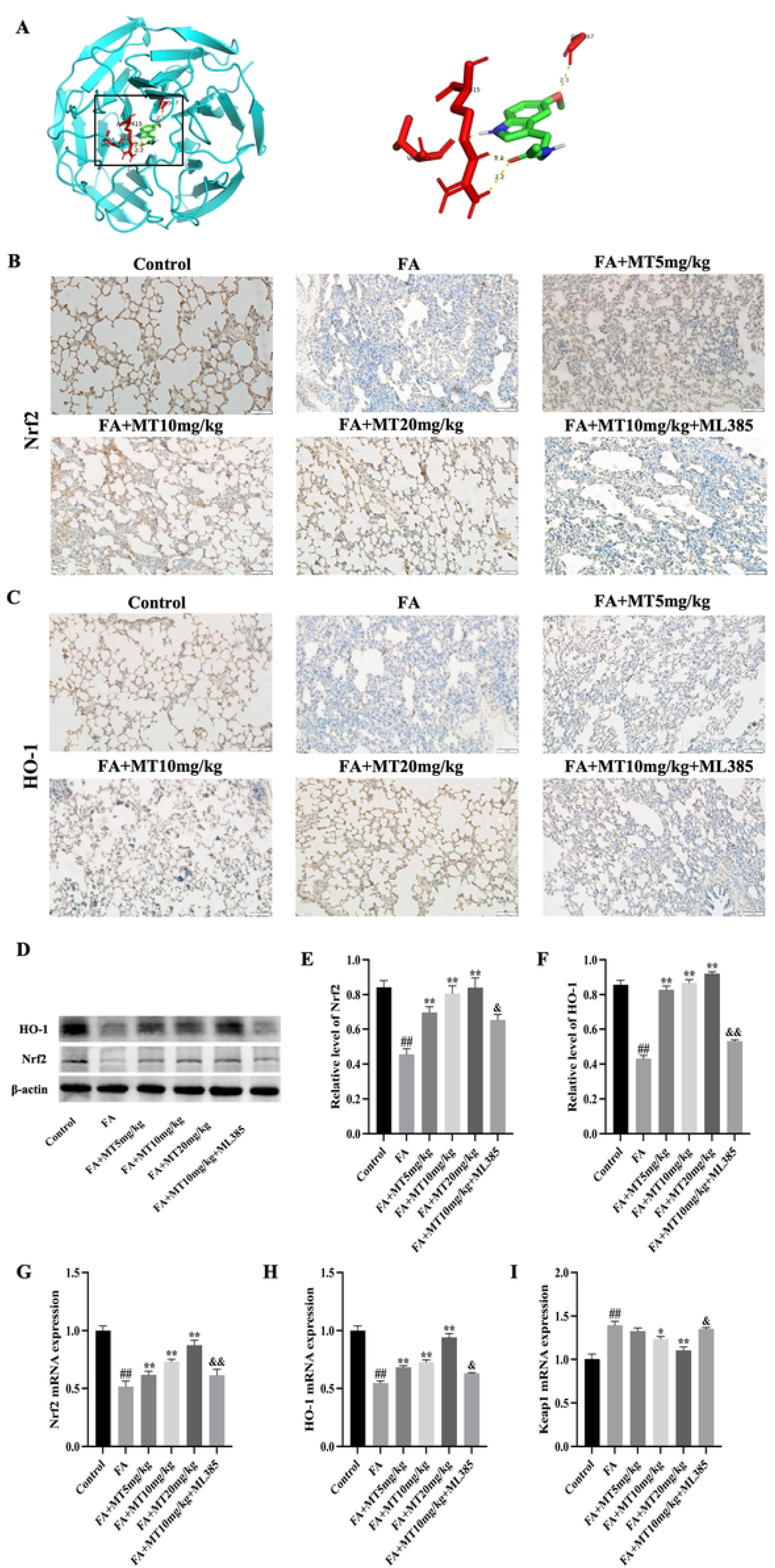
MT antagonizes FA-induced ALI through the Nrf2/HO-1 pathway (A) The docking results of MT with Nrf2. The modelled 3D structure of Nrf2 docked with MT. he enlarged view of binding site in box. Nrf2 protein was shown in color cyan. MT was colored green. The interaction residues showed as color red, bonds showed as yellow dotted lines, and bond lengths were presented as numbers. (B-C) Immunohistochemistry-based images of Nrf2/HO-1 in the lung tissues. Scale bar = 50μm. (D-F) Western blot analysis and quantification of relative grayscale values of Nrf2/HO-1 expression levels for each group. (G) Nrf2 qPCR results. (H) HO-1 qPCR results; (I) Keap1 qPCR results. Animal groups (in all panels): n = 3 rat per group. (*: p < 0.05, **: p < 0.01, compared with the FA group; #: p < 0.05, ##: p < 0.01, compared with the control group. &: p < 0.05).

### 3.4 MT alleviates inflammation and oxidative stress through Nrf2

Western blot analysis showed that the expression level of p-NF-κB in lung tissue of the FA group was higher than that of the control group, which was significantly reduced by MT treatment; however, Nrf2 inhibitor ML385 could reverse the effect of MT (Fig.5A-B). In addition, we also found the changes of inflammatory factors TNF-α, IL-6, IL-1β and oxidation indicators GSH, SOD, 8-OHDG in lung tissue. FA resulted in the decrease of TNF-α, IL-6, and IL-1β products, as well as the decrease of GSH content and SOD activity, and the increase of 8-OHDG content, which is a marker of DNA oxidative damage. Levels of inflammation and oxidative stress were significantly reduced when MT was administered, and similarly the Nrf2 inhibitor ML385 reversed the effects of M (Fig.5C-H). These results suggest that MT alleviates FA-induced lung tissue damage by alleviating inflammatory response and oxidative stress through Nrf2.

**Fig. 5.**
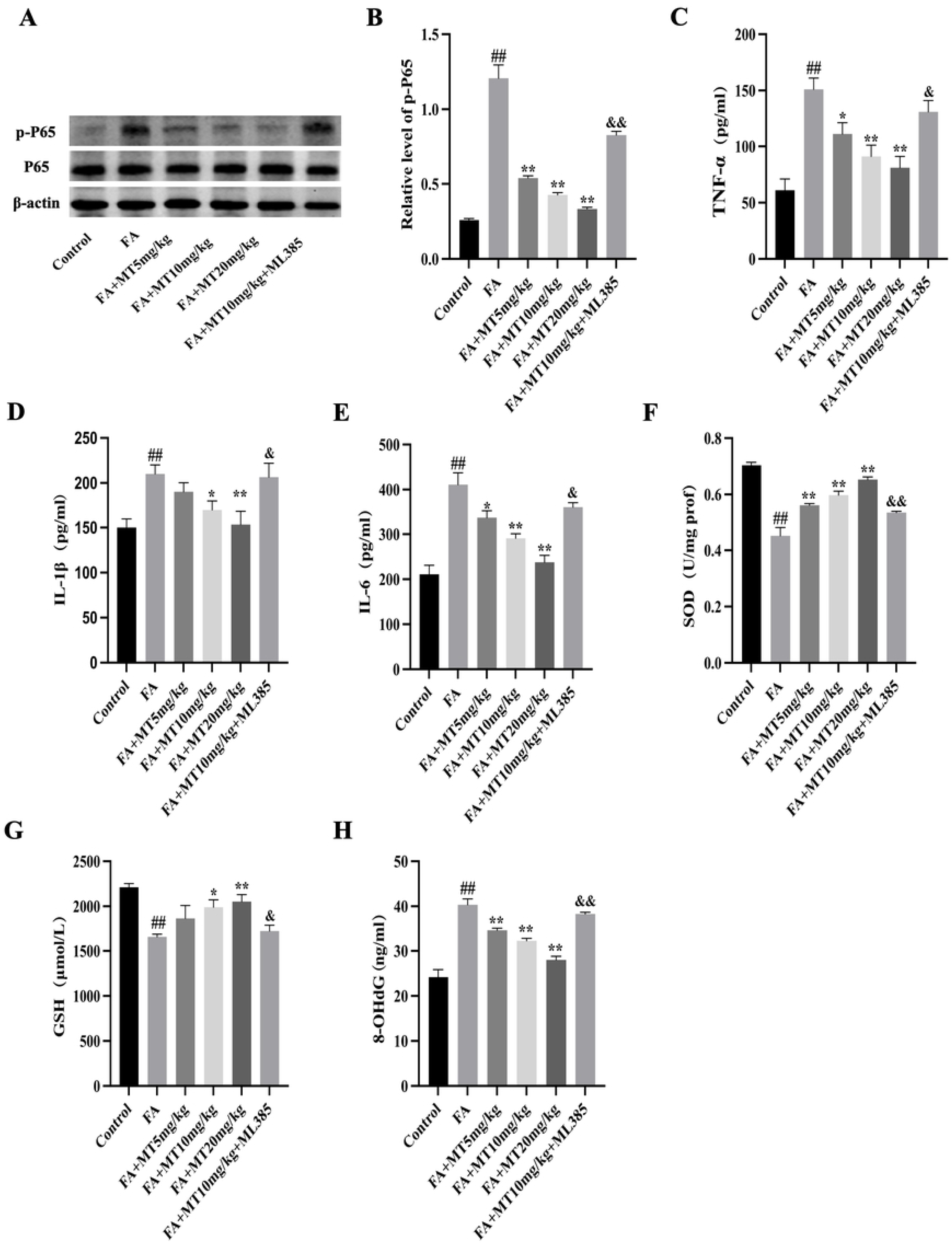
MT alleviates inflammation and oxidative stress through Nrf2 (A-B) Western blot analysis and quantification of relative grayscale values of NFĸB/p-NFĸB expression levels for each group. (C-E) TNF-α, IL-1β and IL-6 concentrations in the BALF. (F-H) Analysis of SOD, GSH and 8-OHdG levels in rat lung tissue by kit. Animal groups (in all panels): n = 3 rat per group. (*: p < 0.05, **: p < 0.01, compared with the FA group; #: p < 0.05, ##: p < 0.01, compared with the control group. &: p < 0.05, &&: p < 0.01, compared with the FA+MT10mg/kg group).

### 3.5 Antagonistic effect of ML385 on the protective effect of MT against acute lung injury

AHR results showed that lung function was improved in the MT group, but decreased after Nrf2 inhibitor administration, which was close to the level of the FA group, suggesting that inhibiting Nrf2 pathway may antagonize the improvement effect of MT on FA-induced lung function abnormalities (Fig.6A). HE staining of lung tissue showed that the degree of lung tissue damage in the Nrf2 inhibitor group was similar to that in the FA group, indicating that Nrf2 inhibitor ML385 could antagonize the protective effect of MT. In addition, lung tissue score, W/D ratio, lung coefficient, Masson and PAS staining further confirmed that MT alleviated FA-induced ALI by activating the Nrf2 pathway (Fig.6B-F).

**Fig. 6.**
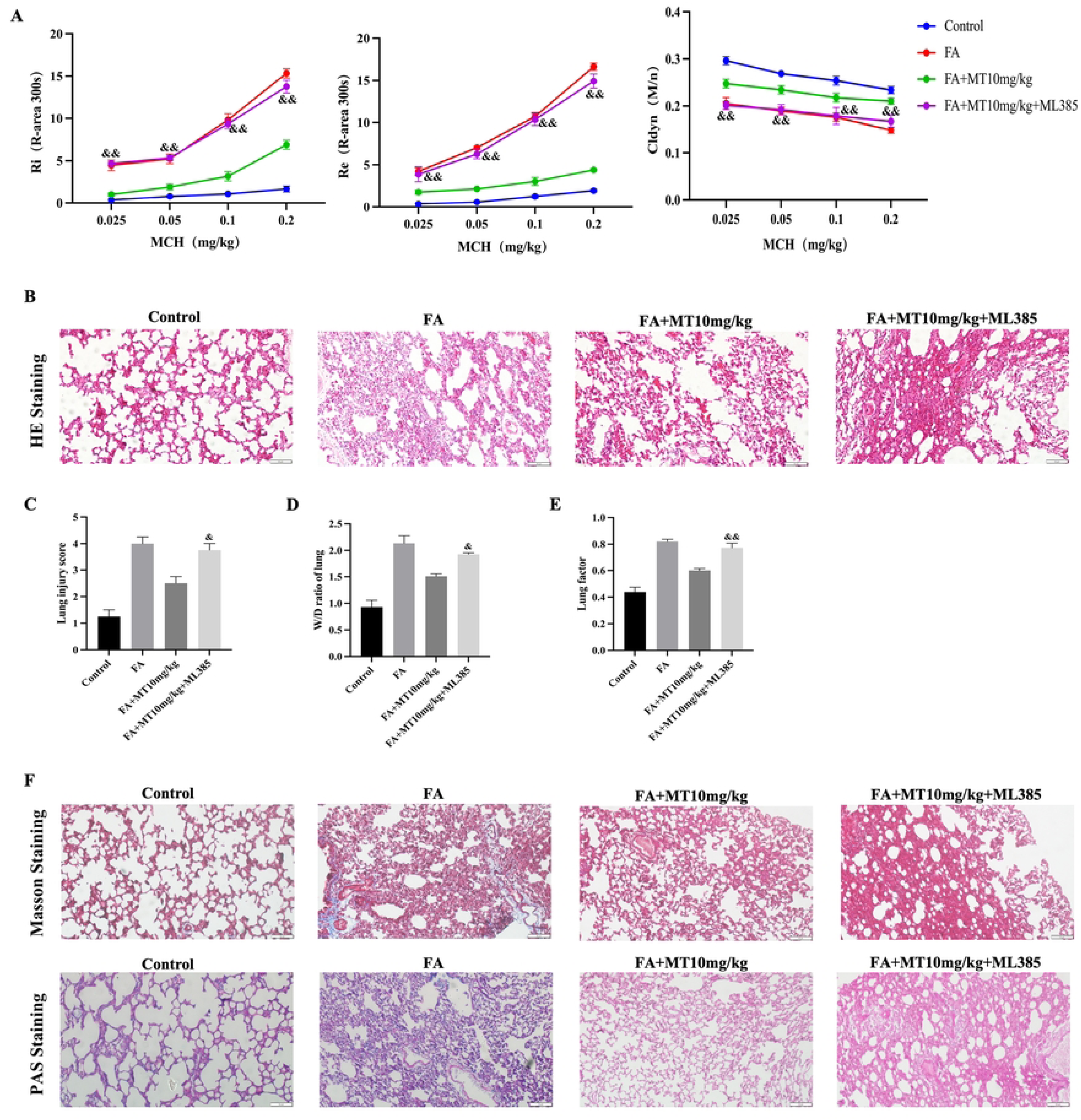
Antagonistic effect of ML385 on the protective effect of MT against acute lung injury (A) Representative H&E staining images of each group of lung tissue sections. Scale bar = 50μm. (B) Quantitative analysis of inflammation score for the H&E staining in each group. (C) Lung wet/dry (W/D) weight ratio. (D) Lung coefficient measurement. (E) Representative Masoon/PAS-stained images of lung tissue sections from various groups. Scale bar = 50μm. (F) R-area of Ri, (G) R-area of Re and (H) peak value of Cldyn (in all panels): n = 3 rat per group. (&&: p < 0.01, compared with the FA+MT0mg/kg group).

## 4. Discussion

ALI/ARDS is an important clinical syndrome associated with high morbidity and mortality, especially in critically ill patients.^21^ The pathological manifestations of ALI primarily involve acute inflammatory responses and dysfunction of alveolar epithelial membrane due to tissue damage.^22^ Moreover, cytokine mediated inflammatory processes aggravate the damage of epithelial and endothelial cells in the pathogenesis of ALI.^23^ FA is a prevalent environmental pollutant found in various sources such as wood products, plastics, synthetic fibers, insulation materials, upholstery, paints, varnishes, household cleaning products, and cigarettes.^24^ Exposure to FA exacerbates the inflammatory response, which is closely associated with its potential mechanism for inducing lung injury. Furthermore, FA exposure can enhance the growth of indoor bacterial communities, and long-term exposure may lead to the development of bacterial communities that pose a high risk to human health.^25^ Although FA is known to cause lung damage, the specific molecular mechanisms underlying this effect remain largely unknown. Therefore, multifaceted validation of new drugs for antagonizing FA-induced ALI is essential.

MT has attracted attention for its various biological activities, such as its anti-inflammatory and antioxidant properties.^27^ However, the potential of MT in the treatment of FA-induced ALI has not been clearly defined. To date, studies have shown that MT has been investigated as a potential treatment for various cross-organ systemic pathological conditions^27^ and that it is also effective in patients infected with neocoronary pneumonia by reducing vascular permeability, anxiety, sedative use and improving sleep quality.^28^ And MT is also a very effective scavenger of superoxide and hydroxyl radicals.^29^ Our study found that this compound also blocks the production of pro-oxidant enzymes by indirectly inhibiting NF-κB. Another indirect antioxidant effect of MT is mediated by Nrf2 transcription factor activation. For example, MT can attenuate diabetes-related restenosis in rats by activating Nrf2 signaling.^30^ Indeed, MT also exhibits critical potential in various respiratory diseases because of the abundance of high-affinity MT receptors captured in lung tissue.^31^

In our study, we discovered that MT played a crucial role in mitigating FA-induced ALI. Through a series of experiments, we observed that MT significantly enhanced lung function. Specifically, it decreased inspiratory and expiratory resistance while increasing lung compliance, thereby reducing airway hyperresponsiveness. This suggests that MT has a positive impact on respiratory function. Additionally, we assessed the structure and function of lung tissue using HE staining, Masson staining, PAS staining, lung injury score, W/D ratio, and lung coefficient. These results indicated that MT reduced inflammatory cell infiltration and collagen fiber proliferation in lung tissue, leading to improved alveolar structure and alleviation of lung injury. Notably, MT decreased the lung injury score and W/D ratio while improving the lung coefficient, indicating its ability to protect the structure and function of lung tissue.

Several studies have shown that Nrf2 is a signaling coordinator that attenuates environmental particulate matter PM2.5-induced lung tissue damage by suppressing inflammation and oxidative stress.^32^ Under normal conditions, Nrf2 binds to Kerch-like ECH-associated protein 1 (Keap1), and when the organism is under oxidative stress, Nrf2 segregates from its negative regulator cytoskeleton-associated protein Kelch-like ECH-associated protein 1 (Keap1) and translocates to the nucleus, where it further promotes transcription of downstream antioxidant genes such as HO-1 upon entry.^33^ To further investigate the role of Nrf2 in MT treatment of FA-induced ALI, we used Nrf2 inhibitor ML385 as an antagonist group. Through IHC, WB and qPCR analysis, we found that ML385 significantly inhibited the increase of Nrf2/HO-1 expression in lung tissue after MT treatment. These results suggest that Nrf2 plays a crucial role in alleviating FA-induced ALI after MT treatment, and provide a valuable reference for further exploring the therapeutic potential of Nrf2 in respiratory diseases.

NF-κB is a nuclear transcription factor that is key signaling molecule of the classical inflammatory pathway that regulates the expression of several genes in the inflammatory response.^34^ Nrf2/Keap1/HO-1 signaling negatively regulates NF-κB transmission in oxidative stress and inflammatory responses, initiating NF-κB-dependent transcriptional pathways that rapidly induce the secretion of inflammatory factors.^35^ Upon activation, NF-κB translocates to the nucleus and binds to specific DNA sequences, leading to the transcriptional expression of inflammatory factors such as IL-1β, TNF-α, and IL-6. This activation also triggers a positive feedback mechanism, further amplifying the inflammatory response. NF-κB plays a crucial role in the body’s immune response and disease progression. It is important to note that the inflammatory response can induce oxidative stress, which further exacerbates inflammation, creating a vicious cycle.^36^ Our study showed that MT reduced expression of inflammatory and increases level of antioxidant in lung tissue, while Nrf2 inhibitor ML385 can reverse this effect, suggesting that MT regulated inflammation and oxidative stress by activating the Nrf2 pathway, thereby alleviating FA-induced ALI.

Studies have shown that FA leads to inflammation closely associated with oxidative stress and the development and progression of ALI. MT, a novel agonist of Nrf2, can resist FA-induced inflammation and oxidative stress, thereby alleviating acute lung injury (Fig.7). This study discloses for the first time that MT supplementation can prevent the deleterious effects of FA on lung tissue, and it is expected that MT could be a possible candidate for the prevention of pollution-induced lung injury.

**Fig. 7.**
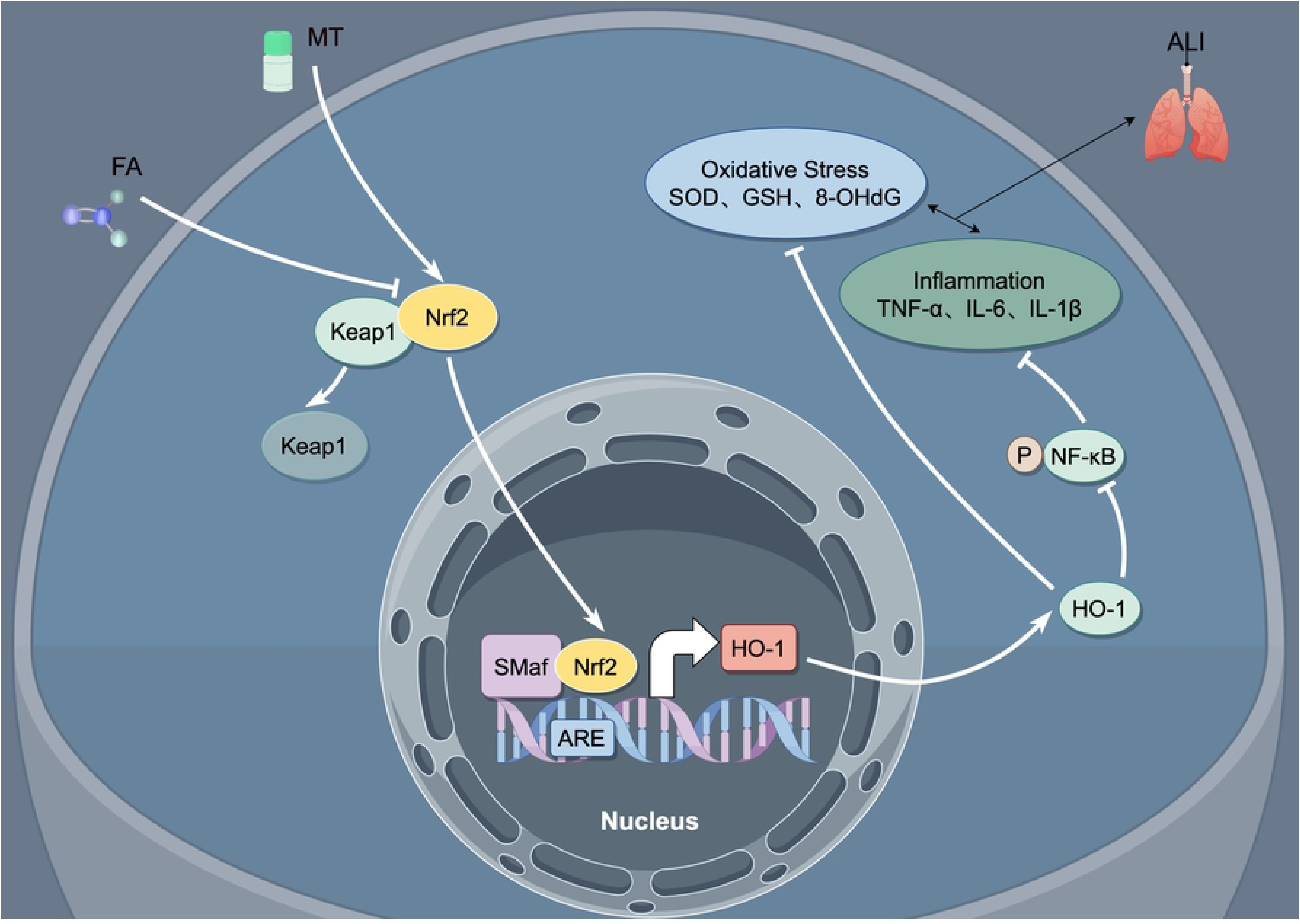
Schematic diagram of the potential mechanisms of MT treatment for acute lung injury caused by FA.

**Funding**

National Natural Science Foundation of China (81902937), Hubei College of Science and Technology, School of Ophthalmology and Stomatology (2020WG06), Hubei College of Science and Technology, School of Ophthalmology and Stomatology (2021WG10).

## References

1. Liu R, Luo X, Li J, Lei Y, Zeng F, Huang X, Yang F. Melatonin: A window into the organ-protective effects of sepsis. J Biomedicine & Pharmacotherapy, 2022, 154, 113556.

2. Störmann P, Becker N, Vollrath J T, Köhler K, Janicova A, Wutzler S, Relja B. Early local inhibition of club cell protein 16 following chest trauma reduces late sepsis-induced acute lung injury. J Journal of Clinical Medicine, 2019, 8 (6), 896.

3. Lai W Y, Wang J W, Huang B T, Lin E P Y, Yang P C. A novel TNF-α-targeting aptamer for TNF-α-mediated acute lung injury and acute liver failure. J Theranostics, 2019, 9 (6), 1741.

4. Zhang X, Zhao Y, Song J, Yang X, Zhang J, Zhang Y, Li R. Differential health effects of constant versus intermittent exposure to formaldehyde in mice: implications for building ventilation strategies. J Environmental science & technology, 2018, 52 (3), 1551–1560.

5. Ding Y, Cui Y, Zhou Z, Hou Y, Pang X, Nie H. Lipopolysaccharide inhibits alpha epithelial sodium channel expression via miR-124-5p in alveolar type 2 epithelial cells. J BioMed Research International, 2020, 2020.

6. Cui Y, Li H, Wu S, Zhao R, Du D, Ding Y, Ji H L. Formaldehyde impairs transepithelial sodium transport. J Scientific reports, 2016, 6 (1), 35857.

7. Chow C W, Herrera Abreu M T, Suzuki T, Downey G P. Oxidative stress and acute lung injury. J American journal of respiratory cell and molecular biology, 2003, 29 (4), 427–431.

8. Vasey C, McBride J, Penta K. Circadian rhythm dysregulation and restoration: the role of melatonin. J Nutrients, 2021, 13 (10), 3480.

9. Wang W, Gao J. Effects of melatonin on protecting against lung injury. J Experimental and therapeutic medicine, 2021, 21 (3), 1–1.

10. Kang J Y, Xu M M, Sun Y, Ding Z X, Wei Y Y, Zhang D W, Fei G H. Melatonin attenuates LPS-induced pyroptosis in acute lung injury by inhibiting NLRP3 GSDMD pathway via activating Nrf2/HO-1 signaling axis. J International Immunopharmacology, 2022, 109, 108782.

11. Arioz B I, Tastan B, Tarakcioglu E, Tufekci K U, Olcum M, Ersoy N, Genc S. Melatonin attenuates LPS-induced acute depressive-like behaviors and microglial NLRP3 inflammasome activation through the SIRT1/Nrf2 pathway. J Frontiers in immunology, 2019, 10: 455021.

12. Gong Y, Yang Y. Activation of Nrf2/AREs-mediated antioxidant signalling, and suppression of profibrotic TGF-β1/Smad3 pathway: a promising therapeutic strategy for hepatic fibrosis—a review. J Life Sciences, 2020, 256, 117909.

13. He F, Ru X, Wen T. NRF2, a transcription factor for stress response and beyond. J International journal of molecular sciences, 2020, 21 (13), 4777.

14. Sajadimajd S, Khazaei M. Oxidative stress and cancer: the role of Nrf2. J Current cancer drug targets, 2018, 18 (6), 538–557.

15. Luo J G, Zhao X L, Xu W C, Zhao X J, Wang J N, Lin X W, Fu Z J. Activation of spinal NF-κB/p65 contributes to peripheral inflammation and hyperalgesia in rat adjuvant-induced arthritis. J Arthritis & Rheumatology, 2014, 66 (4), 896–906.

16. Cao Y, Fan D, Yin Y. Pain mechanism in rheumatoid arthritis: from cytokines to central sensitization. J Mediators of Inflammation, 2020, 2020.

17. Zhao Y, Magaña L C, Cui H, Huang J, McHale C M, Yang X, Zhang L. Formaldehyde-induced hematopoietic stem and progenitor cell toxicity in mouse lung and nose. J Archives of toxicology, 2021, 95, 693–701.

18. Yang Y Q, Ge P, Lv M Q, Yu P F, Liu Z G, Zhang J, Zhou D X. Rno_circRNA_008646 regulates formaldehyde induced lung injury through Rno-miR-224 mediated FOXI1/CFTR axis. J Ecotoxicology and Environmental Safety, 2022, 243, 113999.

19. Wang M L, Wei C H, Wang W D, Wang J S, Zhang J, Wang J J. Melatonin attenuates lung ischaemia–reperfusion injury via inhibition of oxidative stress and inflammation. J Interactive CardioVascular and Thoracic Surgery, 2018, 26 (5), 761–767.

20. Yang B, Ni Y F, Wang W C, Du H Y, Zhang H, Zhang L, Jiang T. Melatonin attenuates intestinal ischemia–reperfusion-induced lung injury in rats by upregulating N-myc downstream-regulated gene 2. J journal of surgical research, 2015, 194 (1), 273–280.

21. Kellner M, Noonepalle S, Lu Q, Srivastava A, Zemskov E, Black S M. ROS signaling in the pathogenesis of acute lung injury (ALI) and acute respiratory distress syndrome (ARDS). J Pulmonary vasculature redox signaling in health and disease, 2017, 105–137.

22. Niethamer T K, Stabler C T, Leach J P, Zepp J A, Morley M P, Babu A, Morrisey E E. Defining the role of pulmonary endothelial cell heterogeneity in the response to acute lung injury. J Elife, 2020, 9, e53072.

23. Kawasaki M, Kuwano K, Hagimoto N, Matsuba T, Kunitake R, Tanaka T, Hara N. Protection from lethal apoptosis in lipopolysaccharide-induced acute lung injury in mice by a caspase inhibitor. J The American journal of pathology, 2000, 157 (2), 597–603.

24. Shao Y, Wang Y, Zhao R, Chen J, Zhang F, Linhardt R J, Zhong W. Biotechnology progress for removal of indoor gaseous formaldehyde. J Applied microbiology and biotechnology, 2020, 104, 3715–3727.

25. Guo J, Xiong Y, Kang T, Zhu H, Yang Q, Qin C. Effect of formaldehyde exposure on bacterial communities in simulating indoor environments. J Scientific Reports, 2021, 11 (1), 20575.

26. Muñoz Jurado A, Escribano B M, Caballero Villarraso J, Galván A, Agüera E, Santamaría A, Túnez I. Melatonin and multiple sclerosis: antioxidant, anti-inflammatory and immunomodulator mechanism of action. J Inflammopharmacology, 2022, 30 (5), 1569–1596.

27. Liu C, Xiao K, Xie L. Advances in the use of exosomes for the treatment of ALI/ARDS. J Frontiers in Immunology, 2022, 4449.

28. Chitimus D M, Popescu M R, Voiculescu S E, Panaitescu A M, Pavel B, Zagrean L, Zagrean A M. Melatonin’s impact on antioxidative and anti-inflammatory reprogramming in homeostasis and disease. J Biomolecules, 2020,10 (9), 1211.

29. Kruk J, Aboul-Enein B H, Duchnik E. Exercise-induced oxidative stress and melatonin supplementation: current evidence. J The Journal of Physiological Sciences, 2021, 71, 1–19.

30. Wang X, Li T, Ma B, Zhao Z, Ni L, Zhang R, Liu C. Melatonin attenuates restenosis after vascular injury in diabetic rats through activation of the Nrf2 signaling pathway. J Biochemical and Biophysical Research Communications, 2021, 548, 127–133.

31. Mazzoccoli G, Kvetnoy I, Mironova E, Yablonskiy P, Sokolovich E, Krylova J, Polyakova V. The melatonergic pathway and its interactions in modulating respiratory system disorders. J Biomedicine & Pharmacotherapy, 2021, 137, 111397.

32. Guohua F, Tieyuan Z, Xinping M, Juan X. Melatonin protects against PM2. 5-induced lung injury by inhibiting ferroptosis of lung epithelial cells in a Nrf2-dependent manner. J Ecotoxicology and Environmental Safety, 2021, 223, 112588.

33. Baird L, Yamamoto M. The molecular mechanisms regulating the KEAP1-NRF2 pathway. J Molecular and cellular biology, 2020, 40 (13), e00099–20.

34. Poma P. NF-κB and Disease. J Int J Mol Sci.2020, 21 (23): 9181.

35. Zhang Q, Liu J, Duan H, Li R, Peng W, Wu C. Activation of Nrf2/HO-1 signaling: An important molecular mechanism of herbal medicine in the treatment of atherosclerosis via the protection of vascular endothelial cells from oxidative stress. J Journal of advanced research, 2021, 34, 43–63.

36. Li J, Deng S H, Li J, Li L, Zhang F, Zou Y, Xu Y. Obacunone alleviates ferroptosis during lipopolysaccharide-induced acute lung injury by upregulating Nrf2 dependent antioxidant responses. J Cellular & Molecular Biology Letters, 2022, 27 (1), 1–20.

